# Evidence for Interaction of 5,10-Methylenetetrahydrofolate Reductase (MTHFR) with Methylenetetrahydrofolate Dehydrogenase (MTHFD1) and General Control Nonderepressible 1 (GCN1)

**DOI:** 10.1101/2024.08.22.609157

**Authors:** Linda R. Büchler, Linnea K.M. Blomgren, Céline Bürer, Vito R.T. Zanotelli, D. Sean Froese

**Affiliations:** Division of Metabolism and Children’s Research Center, University Children’s Hospital Zürich, University of Zürich, Zürich, Switzerland

## Abstract

5,10-methylenetetrahydrofolate reductase (MTHFR) is a folate cycle enzyme required for the intracellular synthesis of methionine. MTHFR was previously shown to be partially phosphorylated at 16 residues, which was abrogated by conversion of threonine 34 to alanine (T34A) or truncation of the first 37 amino acids (i.e. expression of amino acids 38-656), and promoted by methionine supplementation. Here, we over-expressed wild-type MTHFR (MTFHR_WT_), as well as the variants MTHFR_T34A_ and MTHFR_38-656_ in 293T cells to provide further insights into these mechanisms. We demonstrate that following incubation in high methionine conditions (100-1000 *μ*M) MTHFR_WT_ is almost completely phosphorylated, but in methionine restricted conditions (0-10 *μ*M) phosphorylation is reduced, while MTHFR_T34A_ always remains unphosphorylated. Following affinity purification coupled mass spectrometry of an empty vector, MTHFR_WT_, MTHFR_T34A_ and MTHFR_38-656_ in three separate experiments, we identified 134 proteins consistently pulled-down by all three MTHFR protein variants, of which 5 were indicated to be likely true interactors (SAINT prediction threshold of 0.95 and 2 fold-change). Amongst these were the folate cycle enzyme methylenetetrahydrofolate dehydrogenase (MTHFD1) and the amino acid starvation sensor General Control Nonderepressible 1 (GCN1). Immunoprecipitation-immunoblotting of MTHFR_WT_ replicated interaction with both proteins. An AlphaFold 3 generated model of the MTHFR-MTHFD1 interaction places the MTHFD1 dehydrogenase/cyclohydrolase domain in direct contact with the MTHFR catalytic domain, suggesting their interaction may facilitate direct delivery of methylenetetrahydrofolate. Overall, we confirm methionine availability increases MTHFR phosphorylation, and identified potential interaction of MTHFR with MTHFD1 and GCN1.

## Introduction

Methionine is an essential amino acid whose importance to human health is particularly relevant to metabolic diseases, aging and cancer (Parkhitko *et al*., 2019; Bin *et al*., 2024). Intracellular methionine is obtained either from extracellular sources or through recycling via a series of reactions termed the methionine cycle (Lauinger and Kaiser, 2021). This latter cycle constitutes four enzymatic steps, which include (i) the transfer of a folate-derived methyl-group to homocysteine to create methionine, (ii) transfer of methionine to the adenosyl-group of ATP to form S-adenosylmethionine (SAM), (iii) demethylation of SAM to release S-adenosylhomocysteine (SAH), and (iv) breakdown of SAH to adenosine and homocysteine. Both methionine and SAM are important outputs of this cycle. SAM is the most widely used cellular methyl-donor (Cantoni, 1975) and monitoring of its intracellular levels occurs through SAM sensors, including SAMTOR which is part of the mTORC1 complex (Gu *et al*., 2017; Tang *et al*., 2022). Methionine is the initiator amino acid in protein synthesis, and thus methionine availability impacts global protein translation, a mechanism controlled by the amino acid starvation sensor General Control Nonderepressible 1 (GCN1) through its effector protein GCN2 (Tatara *et al*., 2024; Zheng *et al*., 2024).

Integral to methionine production is the enzyme 5,10-methylenetetrahydrofolate reductase (MTHFR). MTHFR generates the folate-derived methyl-group for the creation of methionine by catalyzing the reduction of 5,10-methylenetetrahydrofolate to 5-methyltetrahydrofolate. This reaction is part of the folate cycle, which additionally includes the enzyme methylenetetrahydrofolate dehydrogenase (MTHFD1). Potentially facilitated by the additional role of supplying one carbon units for *de novo* purine synthesis and creation of thymidine, the folate cycle is also intertwined with other core cellular pathways, including the cell cycle (Bar-Joseph *et al*., 2008; Park *et al*., 2015).

MTHFR has been shown to be phosphorylated at 16 residues, most prominently in the first ∼40 amino acids (Froese *et al*., 2018). The phosphorylation priming site appears to be threonine 34, as conversion of this residue to alanine ablates phosphorylation (Yamada *et al*., 2005; Marini *et al*., 2008; Zhu *et al*., 2014). Previous studies have predicted cyclin dependent kinase (CDK) 1 in association with cyclinB1 (Zhu *et al*., 2014), polo-like kinase 1 (Li *et al*., 2017) and DYRK1A/2 (Zheng *et al*., 2019) to be mediators of MTHFR phosphorylation, but confirmatory evidence is lacking. MTHFR phosphorylation sensitizes the enzyme to allosteric inhibition (Froese *et al*., 2018), effected by binding of two SAM molecules to its regulatory domain (Blomgren *et al*., 2024; Yamada, Mendoza and Koutmos, 2024), a state reversed by binding one SAH molecule to the same domain (Froese *et al*., 2018; Blomgren *et al*., 2024; Yamada, Mendoza and Koutmos, 2024). Thus far, little is known about the impact of MTHFR phosphorylation on methionine availability as well as whether it is linked to other SAM or methionine sensing pathways (e.g. translation).

In this work, we have investigated MTHFR in the context of methionine availability and phosphorylation status. We demonstrate that full-length MTHFR over-expressed in 293T cells is preferentially phosphorylated under high methionine conditions. Examination of protein-protein interactions demonstrates no clear differences between phosphorylated full-length wild-type MTHFR, and non-phosphorylated full-length MTHFR harbouring the p.Thr34Ala variant or with an N-terminal truncation of the first 37 amino acids. However, all three MTHFR proteotypes apparently interact with the folate cycle enzyme MTHFD1 and the amino acid sensing protein GCN1. These data expand our knowledge of the metabolic interaction between methionine availability and MTHFR phosphorylation and present novel protein-protein interactions through which these metabolic signals may be affected.

## Results

### Differential phosphorylation in response to methionine availability

We and others have previously demonstrated that full-length wild-type human MTHFR (MTHFR_WT_) expressed in insect (Yamada *et al*., 2005; Froese *et al*., 2018) and human (Zhu *et al*., 2014) cells is phosphorylated at 16 mainly N-terminal residues (**Fig. 1a**). Here, we over-expressed C-terminally Flag-tagged MTHFR_WT_, as well as a variant harbouring the threonine to alanine change at amino acid 34 (MTHFR_T34A_) (**Fig. 1a**), which represents the phosphorylation priming site. These experiments were performed in 293T cells genetically modified to lack endogenous MTHFR activity (**Suppl. Fig. 1a-e**). In standard media conditions, which includes approx. 200 *μ*M methionine (see Methods), immunoblotting with an anti-FLAG antibody detected MTHFR_WT_ as a double band, and MTHFR_T34A_ as a single band, each with a band consistent with degraded protein, which was also present at low levels following expression of an empty vector (EV) (**Fig. 1b**). We expected that the upper band of MTHFR_WT_ represented phosphorylated protein and the lower band unphosphorylated protein, which was confirmed by treatment with alkaline phosphatase, which removed the upper band of MTHFR_WT_ but did not impact MTHFR_T34A_ (**Fig. 1c**).

**Fig. 1.**
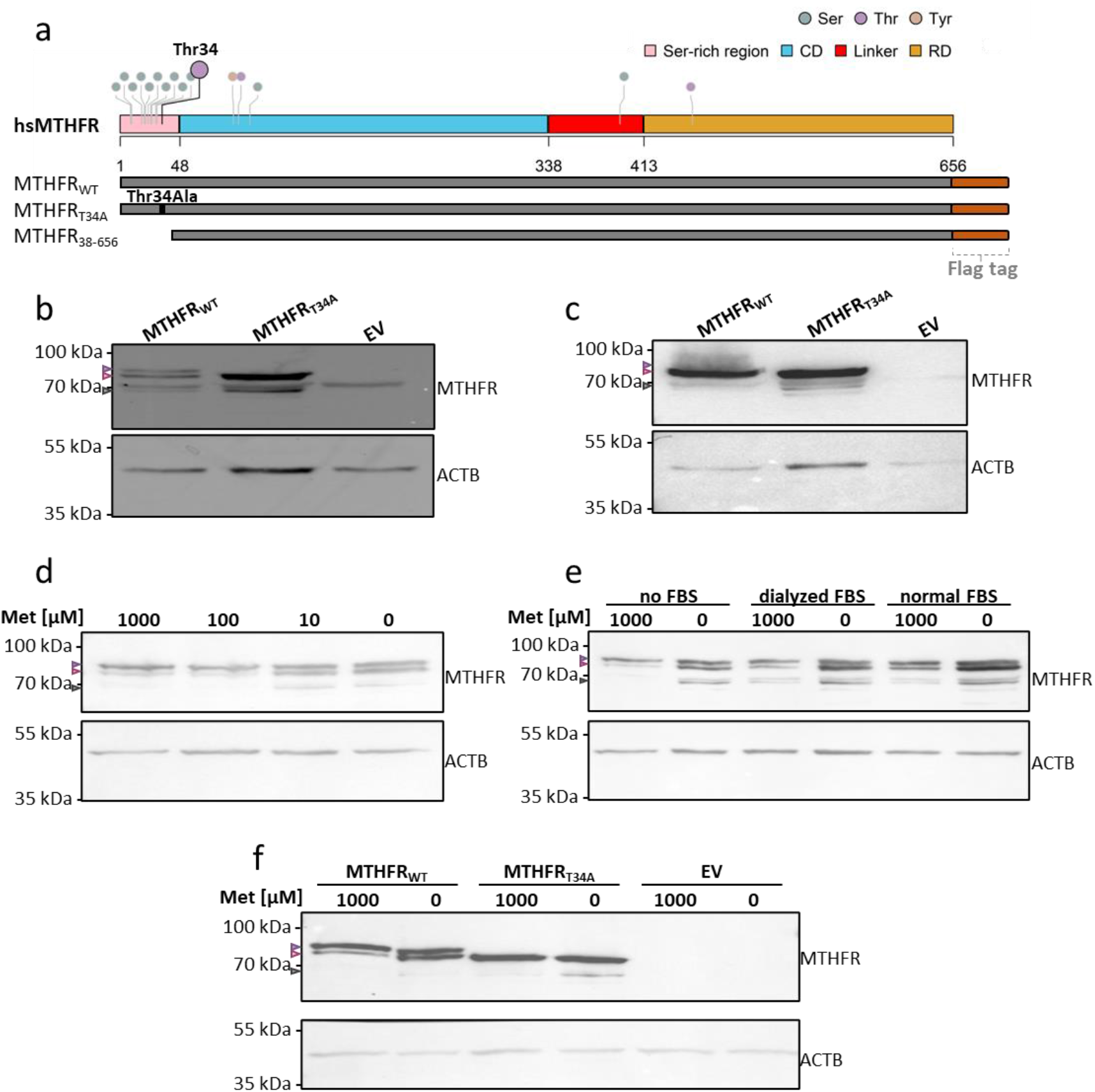
MTHFR phosphorylation status. **a**, Schematic representation of full-length human MTHFR, coloured according to protein domain. Serine rich N-terminal, catalytic domain (CD), linker region and regulatory domain (RD), with the 16 phosphorylation serine, threonine and tyrosine sites highlighted. Below, the generated constructs for expression of MTHFR_WT_, MTHFR_T34A_ and MTHFR_38-656_ with C-terminal FLAG-tag. **b-f**, Over-expression of MTHFR in HEK293T MTHFR knock-out cells visualized using Western blot analysis with the primary antibody targeting the Flag-tag sequence. Expected sizes for MTHFR are marked with triangles for phosphorylated (purple), non-phosphorylated (pink) and degraded protein (gray). Parallel incubation with beta actin (ACTB) was used as loading control, except for in **b**, and **c**, where the membranes were stripped before visualization. Uncropped membranes are available in **Suppl. Fig. 3. b-c**, MTHFR_WT_, MTHFR_T34A_ and empty vector (EV) in standard media conditions containing approx. 200 *μ*M methionine before **b**, and after **c**, alkaline phosphatase treatment. **d-e**, MTHFR_WT_ in media containing **d**, decreasing amounts of methionine (1000 - 0 *μ*M) and **e**, either high concentration (1000 *μ*M) or no (0 *μ*M) methionine, and supplemented with FBS, dialysed FBS or no FBS. **f**, MTHFR_WT_, MTHFR_T34A_ and empty vector (EV) expressed in media with high concentration (1000 *μ*M) or no methionine.

To assess how the availability of methionine impacts MTHFR phosphorylation, we exposed cells harbouring over-expressed MTHFR_WT_ to FBS free media supplemented with increasing concentrations of methionine (0 – 1000 *μ*M) for 24 hours. We found a dose-dependent increase in MTHFR phosphorylation: at 100 *μ*M and 1000 *μ*M methionine, the upper phosphorylated band was the most prominent; while at 0 *μ*M and 10 *μ*M methionine, the intensity of the phosphorylated and non-phosphorylated bands were of approximately equal intensity (**Fig. 1d**). This is consistent with previous results (Zheng *et al*., 2019). Methionine free media containing standard FBS contains low concentrations of methionine (∼2.5 *μ*M), while that containing dialyzed FBS presumably has none. Interestingly, the unphosphorylated band appeared to increase in intensity according to no FBS > dialyzed FBS > normal FBS, in both the presence and absence of methionine (**Fig. 1e** and **Suppl. Fig. 2**). This suggests FBS may contain an additional component which impacts MTHFR expression and phosphorylation. The impact of methionine on MTHFR phosphorylation was supported by replication of the lowest (0 *μ*M) and highest (1000 *μ*M) methionine supplementation conditions in the presence of MTHFR_WT_, MTHFR_T34A_ and EV without visible changes in protein expression (**Fig. 1f**).

### Immunoprecipitation coupled mass spectrometry identifies potentially interacting proteins

Previous studies identified CDK1 (Zhu *et al*., 2014), PLK1 (Li *et al*., 2017) and DYRK1A/2 (Zheng *et al*., 2019) to be potential effectors of MTHFR phosphorylation. We anticipated that performing immunocapture using phosphorylated MTHFR_WT_ as bait, and comparing captured prey proteins against those of non-phosphorylated MTHFR_T34A_ and MTHFR lacking the heavily phosphorylated N-terminal 37 amino acids (i.e. MTHFR_38-656_) may enable identification of the true interacting protein kinase. To this end, we performed three independent immunoprecipitation coupled mass spectrometry experiments using all three MTHFR proteoforms (MTHFR_WT_, MTHFR_T34A_ and MTHFR_38-656_) and EV (**Fig. 2a**). In total, we identified 682 proteins captured by all three MTHFR proteoforms but not EV (**Fig. 2b)**. Only 8 proteins were captured only by MTHFR_WT_ (**Fig. 2b**), but none are annotated as a protein kinase (**Suppl. Table 1**).

**Fig. 2.**
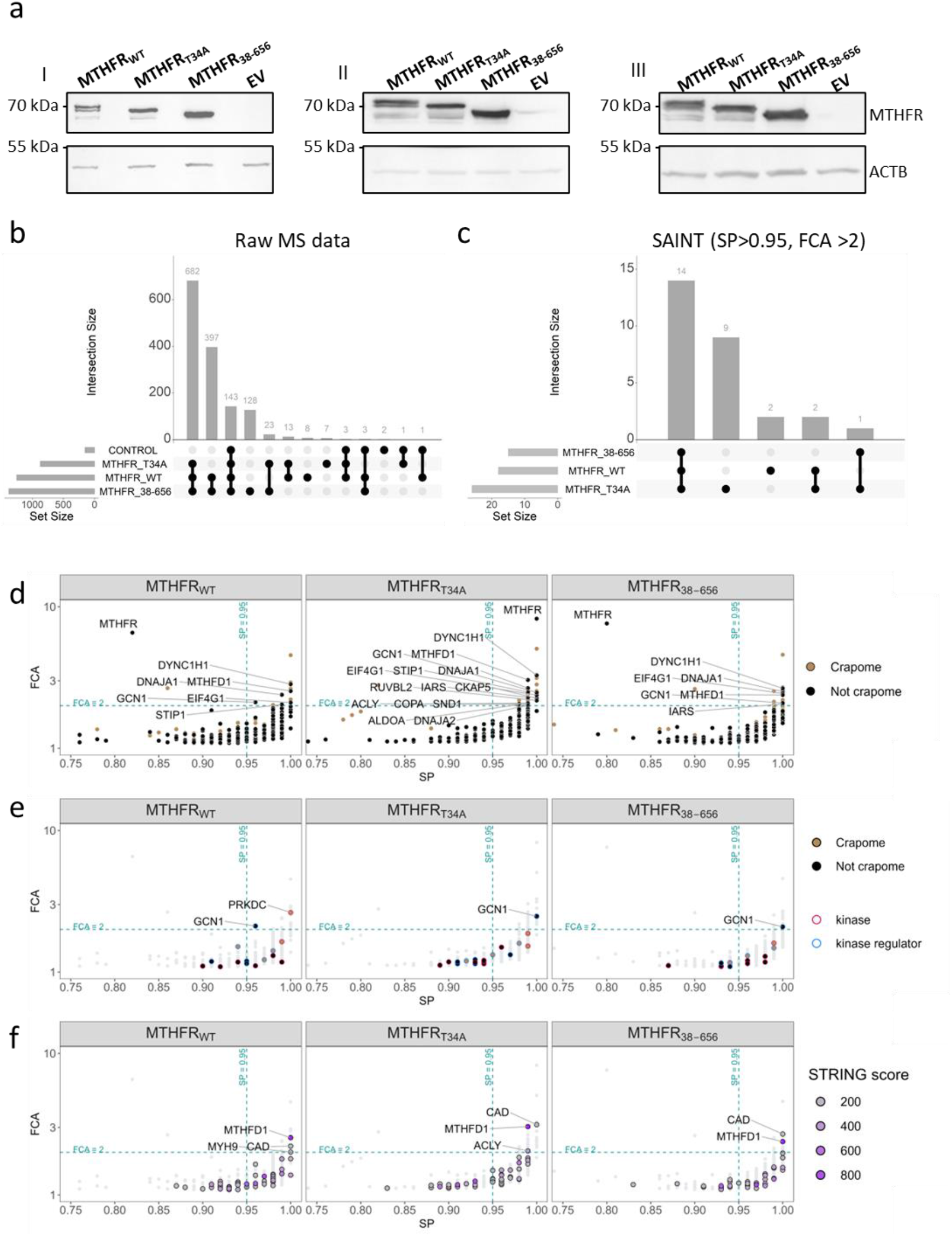
Immunoprecipitation coupled mass spectrometry. Immunoprecipitation followed by LC-MS/MS using constructs MTHFR_WT_, MTHFR_T34A_, MTHFR_38-656_ and empty vector (EV) over-expressed in HEK293T MTHFR knock-out cells. Three biological replicates (I-III) were performed for each construct. **a**, MTHFR in immunoprecipitated samples visualized using Western blot analysis with the primary antibody targeting the Flag-tag sequence. Parallel incubation with beta actin (ACTB) used as loading control. Uncropped membranes are available in **Suppl. Fig. 5. b**, UpSet plots (Conway, Lex and Gehlenborg, 2017) of raw MS data from immunoprecipitation experiments showing overlap in captured protein species between the MTHFR proteoforms and empty vector (EV). **c-f**, SAINT analysis of MS data, using total peptide count and EV as negative control. **c**, UpSet plot showing high probability interactions where SAINT probability (SP) > 0.95 and fold change (FCA) > 2. **d-f**, FCA vs SP graph of all captured proteins, with SP=0.95 and FCA=2 indicated (blue dashed lines). **d**, Protein species present in more than 50% of the experiments in the CRAPome (Mellacheruvu *et al*., 2013) are highlighted in brown. High confidence interactands are labeled. **e**, Proteins with registered protein kinase activity (GO: 0004672) and protein regulator activity (GO: 0019207) are highlighted, high confidence interactands are labeled. **f**, Interactands identified in the STRING database (Szklarczyk *et al*., 2023) with threshold score 100 are highlighted, high confidence interactands are labeled

To discriminate against false positive interactors, we used the SAINT (Significance Analysis of INTeractome) computational tool to probabilistically score the interactions for likely true interactands (Choi *et al*., 2011; Teo *et al*., 2014). A confidence threshold of SAINT probability score of 0.95 and 2-fold change over EV was used to identify likely true interactions. By this method, we identified 14 proteins reliably captured by all three proteoforms but not EV (**Fig. 2c**). As MTHFR was detected at low levels in 2 out of 3 EV replicates (**Suppl. Table 1**), the probabilistic SAINT method scored its likelihood to be a true interactand below the confidence cutoff for MTHFR and MTHFR_38-656_. Using the same cutoffs, no proteins were enriched in MTHFR_WT_ or MTHFR_T34A_ compared to MTHFR_38-656_ (**Suppl. Fig. 4**), suggesting none of the found interactions require the (phosphorylated) N-terminus. Further filtering out of known contaminants by ignoring proteins represented in more than 50% of experiments in the Contaminant Repository for Affinity Purification (CRAPome) (Mellacheruvu *et al*., 2013), identified 6, 14 and 6 proteins confidently captured by MTHFR_WT_, MTHFR_T34A_ and MTHFR_38-656_ respectively (**Fig. 2d**). The confidently captured interactands did not contain the expected putative MTHFR kinases CDK1, PLK1 nor DYRK1A/DYRK2. To specifically identify other interacting kinase or kinase regulators, we filtered our hits using Gene Ontology annotations, identifying two kinase related proteins: GCN1, an activator of the protein kinase GCN2, which was found to interact with all three proteoforms, and PRKDC (protein kinase, DAN-activated, catalytic subunit) identified with MTHFR specifically, but is 57% represented in the CRAPome and thus likely a contaminant (**Fig. 2e**). To identify reported or suggested interactions in our hits, we used the STRING interaction database (Szklarczyk *et al*., 2023). This highlighted the folate cycle protein MTHFD1 as an interactor of all three MTFHR proteoforms (**Fig. 2f**). Of note, the methionine cycle enzyme ACLY was found to interact with MTHFR_T34A_ (**Fig. 2f**). CAD – the pyrimidine biosynthetic enzyme with a reported STRING interaction with MTHFR was found to interact strongly with both MTHFR_T34A_ and MTHFR_38-656_ but is also a known contaminant for AP-MS (CRAPome occurrence 54%). Supported by these analyses, we chose to further validate the MTHFD1 and GCN1 interactions. The full list of all found putative interactions can be found in **Suppl. Table 2**.

### Further evidence of interaction with MTHFD1 and GCN1, modelling of a potential MTHFR-MTHFD1 complex

We used immunoprecipitation-immunoblotting to validate interaction of MTHFR_WT_ with MTHFD1 and GCN1. Using protein specific antibodies, we detected the presence of MTHFD1 and GCN1 in the total cell lysate and after immunoprecipitation with anti-flag following transfection with flag-tagged MTHFR_WT_, but not EV (**Fig. 3a,b** and Suppl. Fig. 6a). Together with the immunoprecipitation coupled mass spectrometry experiments, these data strongly suggest MTHFR physically interacts with both proteins.

**Fig. 3.**
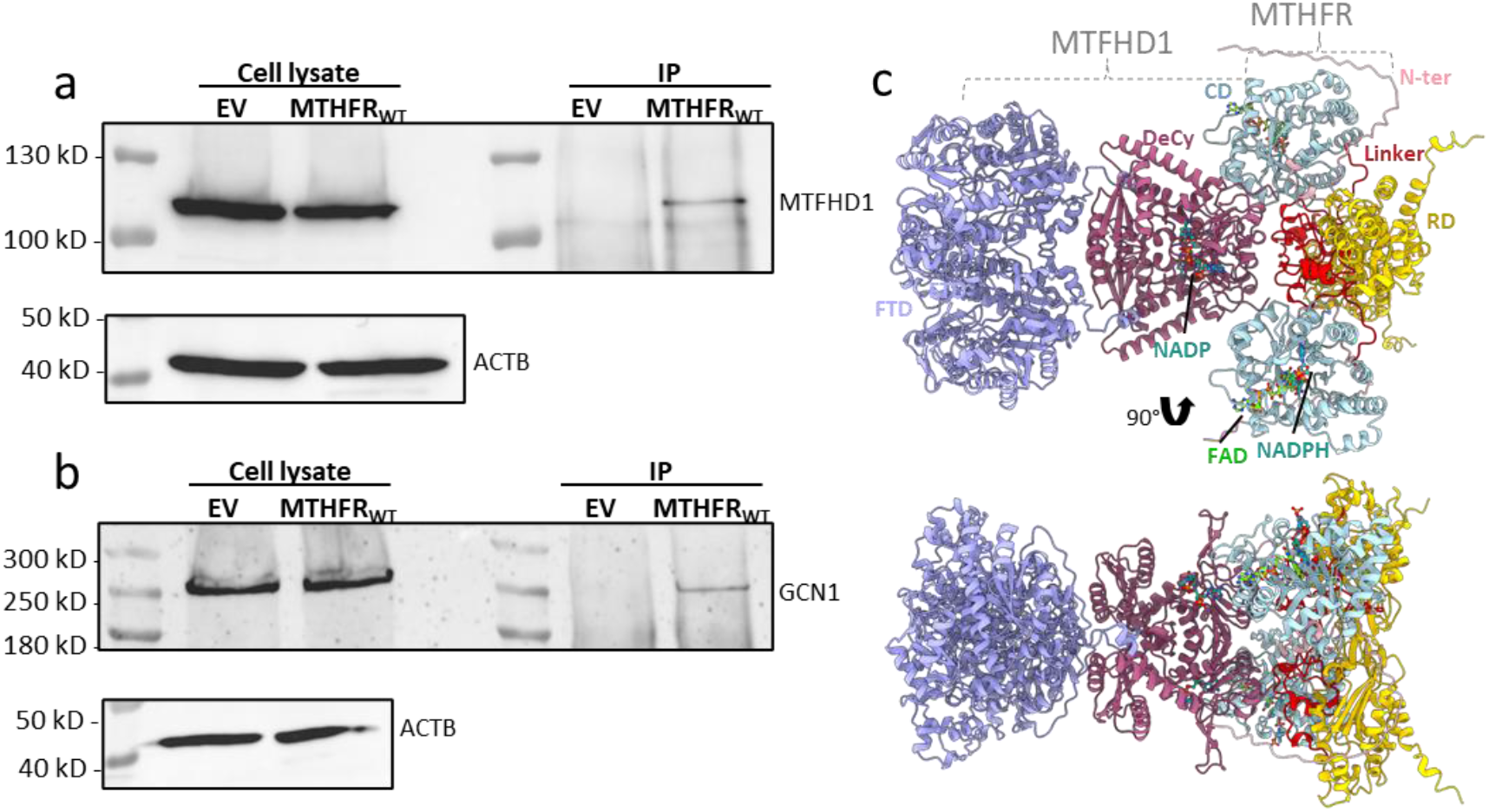
Follow-up analyses support interactions between MTHFR, MTHFD1, and GCN1. Immunoprecipitation of MTHFR_WT_ and empty vector (EV) over-expressed in HEK293T MTHFR knock out cells. Western blot using cell lysate prior to affinity pull down and after immunoprecipitation (IP) visualized with specific primary antibodies targeting **a**, MTHFD1 expected at 101 kDa and **b**, GCN1 expected at 293 kDa. **c**, Alphafold 3 (Abramson *et al*., 2024) prediction of dimeric MTHFR and dimeric MTHFD1 interaction. Colored according to the MTHFD1 dehydrogenase/cyclohydrolase-ligand-domain amino acids 2-291 (DeCy, dark red) and formyltetrahydrofolate synthetase domain amino acids 292-935 (FTD, purple), and the MTHFR catalytic domain (CD, blue with pink N-terminus), linker region (red) and regulatory domain (RD, yellow) for MTHFR. The interaction was predicted with ligands nicotinamide adenine dinucleotide phosphate (NADP) for MTHFD1 and flavine adenine dinucleotide (FAD) and reduced NADPH for MTHFR, which are present in the active sites.

To visualize how MTHFR and MTHFD1 may interact, we created 3D models using AlphaFold 3 (Abramson *et al*., 2024). Although multiple conformations are possible, the highest scored prediction using full-length MTHFR and MTHDF1 as inputs (see Methods) was a conformation in which the dehydrogenase / cyclohydrolase domain of MTHFD1 is positioned adjacent to the catalytic domain of MTHFR (**Suppl. Fig. 6b**,**c**). Prediction made with ligands NADP (MTHFD), FAD (MTHFR) and NADPH (MTHFR) resulted in the same domain orientation except for the position of the flexible N-terminal region (**Fig. 3c)**. In these models, MTHFD1 may transfer the 5,10-methylenetetrahydrofolate created by the dehydrogenase / cyclohydrolase domain directly to MTHFR for further reduction to methyltetrahydrofolate, potentially being an example of substrate channelling (Miles, Rhee and Davies, 1999).

## Discussion

In this study we examined MTHFR phosphorylation as well as potential interacting proteins. Our work is consistent with previous studies that demonstrated a lack of phosphorylation when threonine 34 is mutated to alanine or when the N-terminal serine-rich domain is removed (Yamada *et al*., 2005; Marini *et al*., 2008; Zhu *et al*., 2014; Froese *et al*., 2018), reinforcing the importance of this region to phosphorylation. Likewise, we confirm that methionine availability confers a dose-dependent response to MTHFR phosphorylation. As described previously (Zheng *et al*., 2024), we could confirm that in high methionine conditions MTHFR is predominantly phosphorylated. This can be seen as a form of negative feedback, since phosphorylated MTHFR is sensitized to inhibition, and thus produces less 5-methyltetrahydrofolate, the co-substrate of homocysteine for methionine synthesis. In the absence of supplemented methionine, we detected approximately equal abundances of phosphorylated and non-phosphorylated protein, which is also consistent with the findings of Zheng *et al*.. This suggests that there are methionine-dependent as well as methionine-independent mechanisms to confer phosphorylation, or there is baseline activation of the kinase which confers MTHFR phosphorylation even in conditions with low methionine availability.

Although we found MTHFR to be phosphorylated, we were unable to confirm interaction with the previously described kinases CDK1, DYRK1A and PLK1(Zhu *et al*., 2014; Li *et al*., 2017; Zheng *et al*., 2019), nor did we find strong evidence for direct interaction with any other protein kinase. This does not rule out the participation of the above kinases in MTHFR phosphorylation however. Furthermore, we could not identify any proteins whose interaction depended on the presence of the (phosphorylated) N-terminus of MTHFR, as no proteins were enriched in the full-length MTHFR sample compared to the other two proteoforms. This may have been partially due to the stringency of our cut-offs as we considered only 14 of the 682 proteins found to be associated with all three proteoforms (and not EV) to be likely true, strong interactors. However, we feel this more stringent approach boosts confidence of discovered proteins.

With this latter in mind, we identified a high confidence interaction with GCN1, a kinase related protein that participates in amino acid (including methionine) sensing (Castilho *et al*., 2014). Under amino acid starvation conditions, GCN1 binds ribosomes which have collided due to the accumulation of uncharged tRNAs at ribosomal A-sites (Efeyan, Comb and Sabatini, 2015). Ribosomally-bound GCN1 recruits the kinase GCN2 (Pochopien *et al*., 2021) and initiates the integrated stress response, which among other effects results in a global repression of translation and upregulation of ATF4 which induces genes involved in amino acid biosynthesis (Jin *et al*., 2021). GCN1 is a large protein (2671 amino acids, NP_006827.1) composed mainly of HEAT repeats, which typically function as protein-protein interaction surfaces (Andrade and Bork, 1995). This corresponds well to the over 350 proteins suggested to interact with GCN1 (NCBI, Gene: 10985, updated 17 June 2024), and makes unclear whether MTHFR directly interacts with GCN1 or through an intermediary protein.

On the other hand, MTFHD1 represents the preceding step in folate metabolism to MTHFR, and it would be expected that their interaction is direct. Thus far little is known about mechanisms driving the fate of the MTHFD1 product 5,10-methylenetetrahydrofolate, which is a substrate of thymidylate synthase (TYMS) for the creation of thymidine 5’-monophosphate, a substrate of serine hydroxymethyltransferase (SHMT1) to create serine from glycine, and a substrate of MTHFR for reduction to 5-methyltetrahydrofolate (Tibbetts and Appling, 2010). It may be surmised that a direct enzymatic interaction between MTHFD1 and MTHFR is part of a fate determining process for 5,10-methylenetetrahydrofolate. The Alphafold 3 model, which places the 5,10-methylenetetrahydrofolate producing cyclohydrolase/dehydrogenase domain of MTHFD1 in direct contact with the 5,10-methylenetetrahydrofolate utilizing catalytic domain of MTHFR would be consistent with such a role. However, caution should be used in interpreting such *in silico* models without experimental evidence. Nevertheless, it would be interesting to determine whether MTHFD1 additionally interacts with TYMS and SHMT1.

This study benefitted from the implementation of three MTHFR proteoforms, including phosphorylated and non-phosphorylated variants, expressed in a cell line that had no endogenous MTHFR. Our methodology enabled comparison across proteoforms as well as replicates, allowing a stringent determination of interacting proteins using a data-driven approach for which it can often be challenging to identify true interactors from background contaminants (Mellacheruvu *et al*., 2013). For this reason, it was important that we were able to replicate identification of our two most interesting target proteins using immunoprecipitation-immunoblotting. Nevertheless, our approach and subsequent interpretations remain limited by the requirement of over-expression, the fact that our negative control (EV) did not account for interactions accrued due to protein over-expression, that we did not have a fully phosphorylated proteoform, and that all interaction studies were antibody-based. Further, we found trace levels of MTHFR in two of our negative control (EV) immunoprecipitation samples, consistent with detection of a faint band of flag-tagged MTHFR in Western blots, which decreased our power to identify weak protein-protein interactions. For future studies, it would be very interesting to determine whether methionine or amino acid availability impacts interaction of MTHFR and GCN1, and likewise if folate availability impacts interaction of MTHFR and MTHFD1. Additionally, it would be valuable to map whether these interactions are direct or indirect, and any impact their disruption may have on core cellular processes.

## Methods

### Cloning

Modified pcDNA3-c-flag-LIC (#20011, Addgene), a generous gift from SGC, was used as the backbone in cloning experiments, allowing for expression with a C-terminal fusion sequence SSKGGYGSDYKDDDDK (Flag-tag). The construct MTHFR_WT_ was generated by cloning the human wild-type MTHFR (IMAGE: 6374885) sequence into the backbone (Weile *et al*., 2021), as well as the same sequence carrying the point mutation threonine to alanine at position 34 (MTHFR_T34A_). Primers can be found in **Suppl. Table 3**.

Ligation-independent cloning (LIC) was employed for constructing truncated MTHFR_38-656_ using MTHFR_WT_ as gene template. The gene insert was amplified using touchdown PCR (Korbie and Mattick, 2008) and NEB Phusion DNA polymerase (#M0530L, NEB). Primer sets for obtaining the gene insert can be found in **Suppl. Table 3**. The insert was digested with DpnI (#R0176L, NEB) and annealed into BsaI (#R3435S, NEB) digested pcDNA3-c-flag-LIC (#20011, Addgene) backbone by incubating for 15 minutes at room temperature, then transferring to ice.

### Tissue culture and transfection

HEK293T (CRL-3216, ATCC) MTHFR knock-out cells were generated by CRISPR-Cas9 base editing (Ran *et al*., 2013) (guide RNA found in **Suppl. Table 3)**, whereby the modification c.517_525delins91 was introduced by non-homologous end joining, resulting in skipping of exon 4 and no enzymatic activity (**Suppl. Fig. 1**). Enzymatic activity was determined using an HPLC-based assay originally described by Suormala et al. (Suormala, Gamse and Fowler, 2002) with previously described adaptations (Burda *et al*., 2015, 2017). Cells were cultured in DMEM with Glutamax (#31966-021, ThermoFisher) containing 10% FBS (#10270-106, Gibco) and 1% antibiotic/antimycotic (#15240-062, Gibco). At 60-90% cell confluency, the cells were transfected with Lipofectamine 3000 (#L3000-008, Invitrogen), using constructs MTHFR_WT_, MTHFR_T34A_ and MTHFR_38-656_ in overexpression experiments. Unmodified pcDNA3-c-flag-LIC (#20011, Addgene) was used as an empty vector control throughout all experiments. GFP was used as a control of transfection efficiency. After 24 hours of incubation, the cells were either subjected to continued cultivation in methionine concentration experiments or after a total of 48 hours following transfection, cells were washed with PBS before being harvested for western blotting or affinity pull-down experiments.

### Methionine concentration experiments

Transfections of MTHFR_WT_, MTHFR_T34A_, and empty vector (EV) were, 24 hours after transfection, washed with PBS and incubated again for 24 hours in media containing high glucose, no glutamine, no methionine, and no cystine (#21013-024, Gibco). The media was supplemented with either 1000 *μ*M, 100 *μ*M, 10 *μ*M, or 0 *μ*M methionine (#M5308, Sigma-Aldrich; filter sterilised) and 0.2 mM cysteine (#C7352, Sigma-Aldrich; filter sterilised), 1:100 GlutaMAX (#35050-038, Gibco), and 1:100 antibiotic/antimycotic (#15240-062, Gibco). Additionally, the media contained either no FBS, 10% V/V FBS (#10270-106, Gibco), or dialyzed FBS (#A33820-01, Gibco). Cells were harvested and used for western blotting and affinity pull-down experiments. Alkaline phosphatase treatment was performed over night at 4°C using Lambda Protein Phosphatase (#P0753S, BioLabs).

### Immunoprecipitation

Harvested cells were crosslinked using 0.5% paraformaldehyde for 10 minutes at room temperature, then quenched with 19 mM glycine for 8 minutes on ice. The cells were centrifuged at 2000 rcf, 4°C, for 5 minutes. The supernatant was removed, and the pellet was suspended in IP lysis buffer (150 mM NaCl, 50 mM Tris pH 8.0, 1% NP-40 (IGEPAL CA-630), 0.5% sodium deoxycholate) containing protease and phosphatase inhibitors (#1861281, Thermo Scientific). The lysates were centrifuged at 10,000 rcf, 4°C, for 5 minutes and protein concentration was determined using the Bradford method. 500 *μ*g of protein in IP lysis buffer was incubated for 10 minutes rotating at room temperature with Dynabeads covalently coupled to Protein G (#10004D, Invitrogen), capturing anti-Flag antibody (#F3165, Sigma). The beads were then washed 3 times (respectively 4 times if samples were used for LC-MS/MS) in PBS and stored in 100*μ*l PBS. Beads were either used in further western blotting experiments or analysed using LC-MS/MS.

### Western Blotting

Western blotting analysis were performed using hand-casted SDS-PAGE gels with a 5% stacking gel and a 7.5%, 10% or 14% running gel. All blots were performed using nitrocellulose membranes (#10600007, Cytiva) with a semi-dry transfer before being blocked with 5% milk TBS-Tween (0.1-0.2% Tween), followed by incubation with the corresponding antibodies development with either chemiluminescence detection reagents Clarity Normal (#170-5060, Bio-Rad) or Clarity Max (#1705062, Bio-rad) and imaging with a ChemiDoc Touch Imaging System (Bio-Rad). Uncropped images are available in **Suppl. Fig. 3** and **Suppl. Fig. 5**. To target MTHFR_WT_, MTHFR_T34A_, MTHFR_38-656_, or EV, a 1:2000 dilution of anti-Flag mouse antibody (#F3165-1MG, Sigma) was used as the primary antibody. For the secondary antibody, a 1:5000 dilution of mouse IgG kappa binding protein conjugated to Horseradish Peroxidase (#sc-516102, Santa Cruz) was used. To target MTHFD1, a 1:2000 dilution of anti-MTHFD1 rabbit antibody (#10794-1-AP, Proteintech) was used. For GCN1, a 1:2000 dilution of anti-GCN1L1 rabbit antibody (#A301-843A, Bethyl) was used. The secondary antibody for these was a 1:1000 dilution of HRP-conjugated anti-rabbit antibody (#sc-2357, Santa Cruz) or a 1:5000 dilution of HRP-conjugated anti-rabbit antibody (#sc-2301, Santa Cruz). As a loading control, β-Actin was used and incubated in parallel with the rest of the membrane using a 1:5000 dilution of monoclonal mouse anti-β-Actin antibody (#A1978, Sigma). The secondary antibody was a 1:5000 dilution of HRP-conjugated anti-mouse antibody (#sc-516102, Santa Cruz). Where noted, ACTB was visualized after mild stripping of the initial blot for 20 minutes at room temperature and re-blocked overnight in 5% milk TBS-T at 4°C before being incubated with the antibodies.

### Mass spectrometry

LC-MS/MS was performed at Functional Genomics Center Zurich (FGCZ). Dynabeads from immunoprecipitation experiments were washed once with 100 *μ*l of 10 mM Tris, 2 mM CaCl_2_, pH 8.2 before digestion at 60°C for 30 minutes in digestion buffer (45 *μ*l of 10 mM Tris, 2 mM CaCl_2_, pH 8.2; 5 *μ*l trypsin (100 ng/*μ*l in 10 mM HCl) and 0.3 *μ*l 1M Tris pH 8.2. Peptides were extracted from the beads with 150 *μ*l of 0.1% trifluoroacetic acid containing 50% acetonitrile. The supernatants were combined and dried. Before analysis, the samples were dissolved in 20 *μ*l ddH2O with 0.1% formic acid and then injected into a nanoAcquity UPLC coupled to a Q-Exactive mass spectrometer (Thermo).

## Data analysis

Mascot (Matrix Science, London, UK; version 2.6.2 and 2.7.0.1) was used to interpret raw MS data. Mascot was set up to search against a UniProt homo sapiens reference (taxonomy: 9606, canonical sequences, 20416 forward sequences, retrieved 2019-07-09), extended with its reversed database as a decoy and 296 contaminants and assuming the digestion enzyme trypsin. Mascot searches were performed with a fragment ion mass tolerance of 0,030 Da and a parent ion tolerance of 10,0 PPM. Oxidation of methionine was specified as a variable modification. Scaffold (version Scaffold_5.3.3, Proteome Software Inc., Portland, OR) was

used to validate MS/MS based peptide and protein identifications. Peptide identifications were accepted if they could be established at greater than 5.0% probability to achieve an FDR less than 0.1% by the Peptide Prophet algorithm (Keller *et al*., 2002) with Scaffold delta-mass correction. Protein identifications were accepted if they could be established at greater than 22.0% probability to achieve an FDR less than 1.0% and contained at least 2 identified peptides. Protein probabilities were assigned by the Protein Prophet algorithm (Nesvizhskii *et al*., 2003). Proteins that contained similar peptides and could not be differentiated based on MS/MS analysis alone were grouped to satisfy the principles of parsimony. Proteins sharing significant peptide evidence were grouped into clusters. SAINT analysis (Choi *et al*., 2011; Teo *et al*., 2014) was performed via https://reprint-apms.org/ with default setting and without CRAPome controls (Mellacheruvu *et al*., 2013), instead CRAPome data was manually downloaded from the CRAPome repository (date: 31/05/2024). Proteins interacting with MTHFR showing up over 50% within the dataset were determined to be a CRAPome. Human protein with kinase activity (GO: 0004672, date: 26/07/2024) and kinase regulator activity (GO:0019207, date: 03/06/2024) were retrieved from amigo.geneontology.org. String data was obtained using R library STRINGdb (Szklarczyk *et al*., 2023) (date: 30/05/2024, v12.0). R scripts for analysis of raw MS data and SAINT output can be found in GitHub “MTHFR and Friends “. Structure prediction and interactions of MTHFR and MTHFD1 were generated with Alphafold 3 and visualised in ChimeraX (Meng *et al*., 2023). Prediction was made without and with ligands FAD and NADPH (MTHFR) and NADP (MTHFD1), generated with seed: 2027060258. Sequences used are provided in **Suppl. Fig. 6b**.

## Data availability

R script are available at GitHub (https://github.com/FroeseLab/mthfr-and-friends)

## Acknowledgements

We thank the Functional Genomics Centre of the University of Zurich who performed digestion of sample beads, run of LC-MS/MS and analysis of data using Mascot. Funding for this study was provided by the EMDO Stiftung and the Theodor und Ida Herzog-Egli Stiftung to L.R.B.. D.S.F. is supported by the Swiss National Science Foundation [310030_192505 and 320030E_219127] and the University Research Priority program of the University of Zurich ITNERARE – Innovative Therapies in Rare Diseases.

## Author contributions

Conceptualization and funding procurement was carried out by L.R.B. and D.S.F. The methodology was the responsibility of L.R.B.. Investigation/data analysis was carried out by L.R.B., L.K.M.B., and V.R.T.Z.. Writing of the manuscript was performed by L.R.B., L.K.M.B., and D.S.F. with contributions and editing from all authors.

## Competing interests

All authors declare no competing interests.

**Correspondence and requests for materials** should be addressed to D. Sean Froese

