## Supplementary Figures and Table 3 for "Evidence for Interaction of 5,10-Methylenetetrahydrofolate Reductase (MTHFR) with Methylenetetrahydrofolate Dehydrogenase (MTHFD1) and General Control Nonderepressible 1 (GCN1)"

### Supplementary data

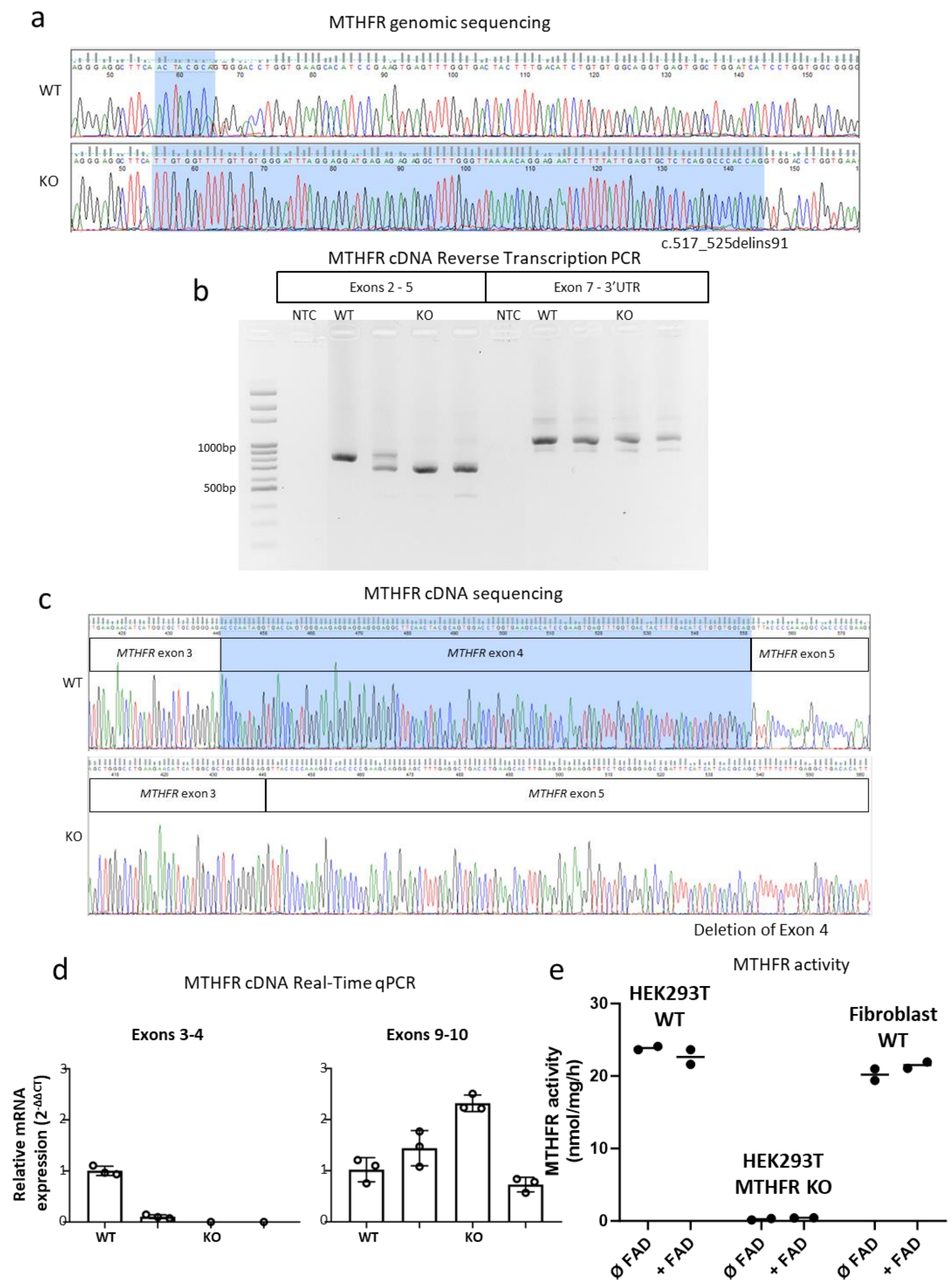


**Supplementary Fig. 1 | Characterisation of generated HEK293T MTHFR knock-out cells (KO) compared to wild-type (WT)**

**a,** Genomic sequencing with highlighted gene insert in KO compared WT **b,** Gel electrophoresis with reverse transcription PCR for WT, KO and no template control (NTC) for exons 2-5 and exons 7- 3´UTR. **c,** Sequence of cDNA covering exons 3,4 and 5 for WT and 3 and 5 for KO. **d,** Real-time qPCR for exons 3-4 and exons 9-10. **e,** Activity for cell-lysate of wild-type HEK293T cells, HEK293T MTHFR knock-out cells and control wild-type fibroblast cell line with and without supplemented FAD.

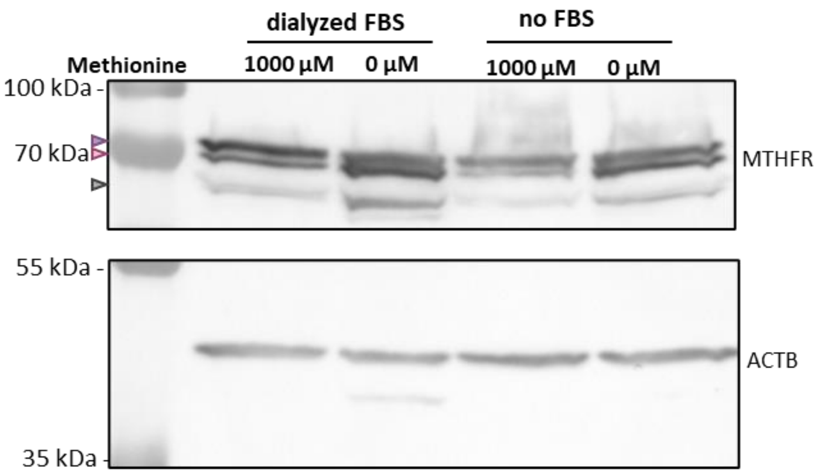


**Supplementary Fig. 2 | MTHFR phosphorylation status**

Over-expressed $MTHFR_{WT}$ in HEK293T MTHFR knock-out cells in media containing either high concentration (1000 µM) or no methionine (0 µM), supplemented with dialysed FBS or no FBS. Visualized using Western blot analysis with the primary antibody targeting the Flag-tag sequence. Expected sizes for MTHFR are marked with triangles for phosphorylated (purple), non-phosphorylated (pink) and degraded protein (gray). Parallel incubation with beta actin (ACTB) was used as loading control.

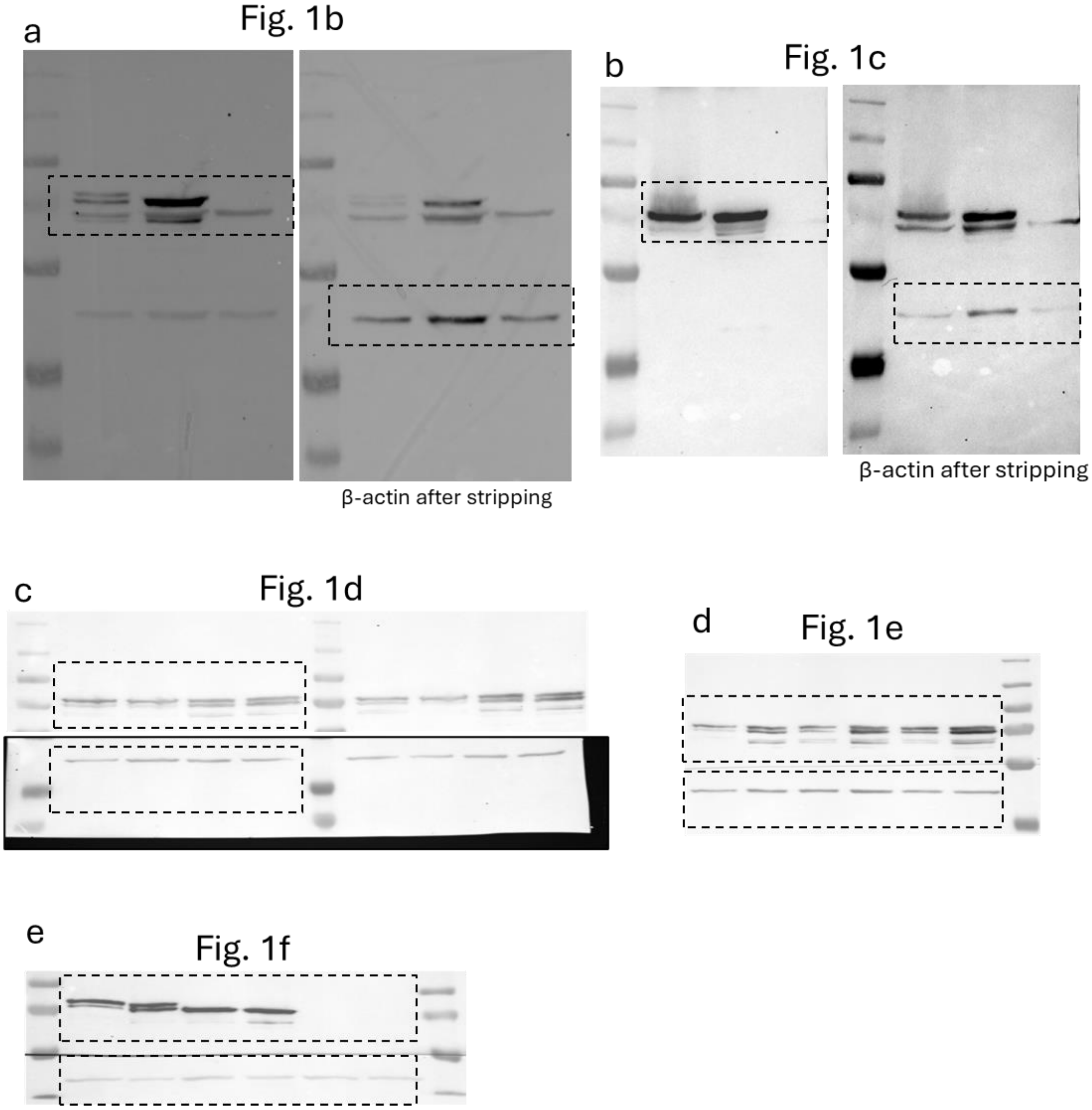


**Supplementary Fig. 3 | Uncut Western Blots from Fig 1**

Shown are the uncut membranes of the Western blotting analysis presented in Figure 1. Dashed rectangle represents the area shown in Figure 1. When different exposures were used for the anti-flag and anti-ACTB antibodies, both exposures are shown.

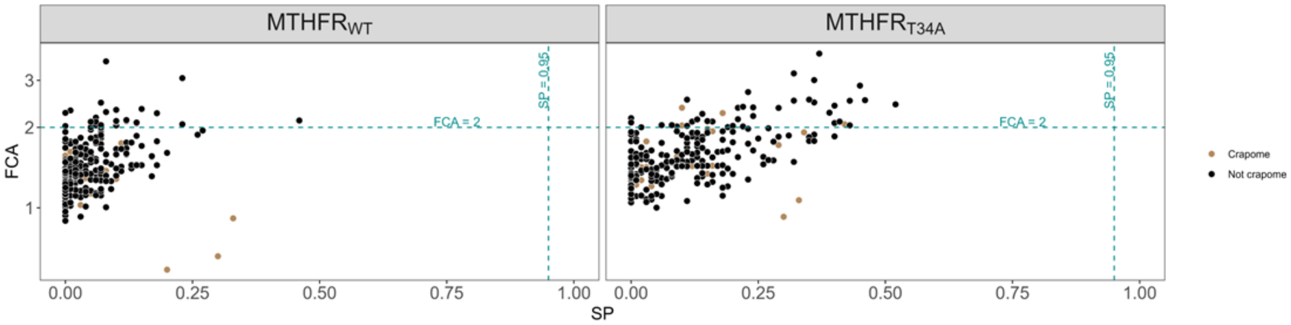


**Supplementary Fig. 4 | Immunoprecipitation coupled mass spectrometry**

FCA vs SP graph of all captured proteins with SP=0.95 and FCA=2 threshold marked, using MTHFR$_{38\text{-}656}$ as negative control. Protein species present in more than 50% of the experiments in the CRAPome (Mellacheruvu *et al.*, 2013) are highlighted in brown.

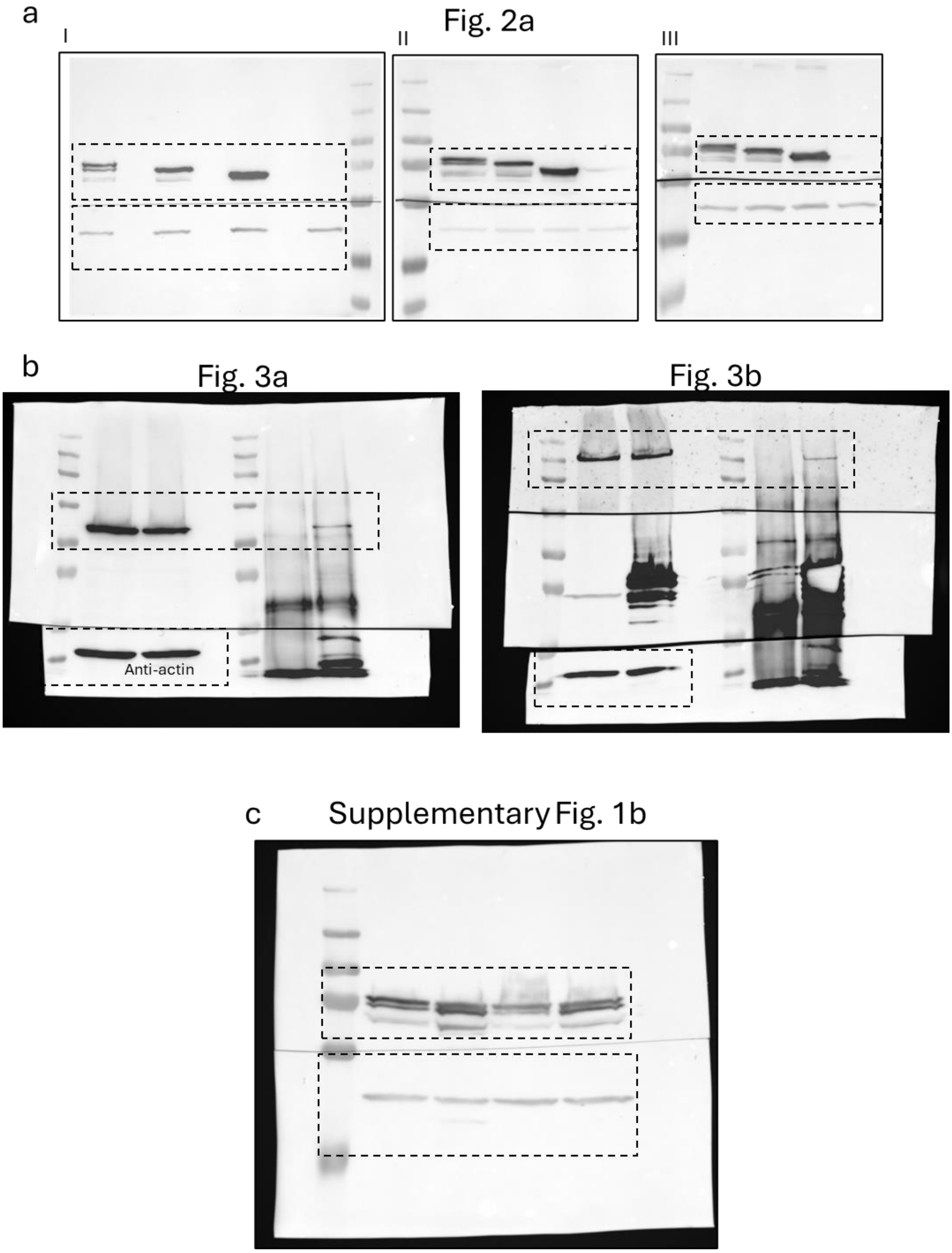


**Supplementary Fig. 5 | Uncut Western blots from Fig 2-3, Supplementary Fig. 1b**
Shown are the uncut membranes of the Western blotting analysis presented in Figures 2, 3 and Supplementary Figure 1b. Dashed rectangle represents the area shown in the other Figures.

a

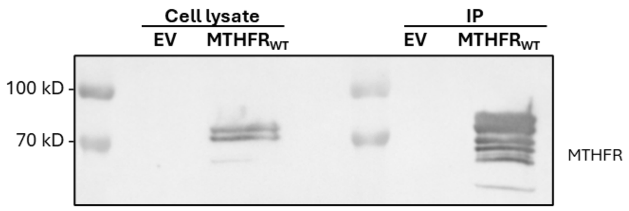


b

**Alphafold3 protein sequence**

Human MTHFR_WT (IMAGE: 6374885)

MVNEARGNSSLNPCLEGSASSGSESSKDSSRCSTPGLDPERHERLREKMRRRLESGDKWFSLEFFPPRTAEGAVNLISRFDRMAAGG
PLYIDVTWHPAGDPGSDKETSSMMIASTAVNYCGLETILHMTCCRQRLEEITGHLHKAKQLGLKNIMALRGDPIGDQWEEEEGGFN
YAVDLVKHIRSEFGDYFDICVAGYPKGHPEAGSFEADLKHLKEKVSAGADFIITQLFFEADTFFRFVKACTDMGITCPIVPGIFPIQGYHS
LRQLVKLSKLEVPQEIKDVIEPIKDNDAAIRNYGIELAVSLCQELLASGLVPGLHFYTLNREMATTEVLKRLGMWTEDPRRPLPWALSA
HPKRREEDVRPIFWASRPKSYIYRTQEWDEFPNGRWGNSSSPAFGELKDYYLFYLKSKSPKEELLKMWGEELTSEASVFEVFVLYLSGE
PNRNGHKVTCLPWNDEPLAAETSLLKEELLRVNRQGILTINSQPNINGKPSSDPIVGWGPSGGYVFQKAYLEFFTSRETAEALLQVLKK
YELRVNYHLVNVKGENITNAPELQPNAVTWGIFPGREIIQPTVVDPVSFMFWKDEAFALWIEQWGKLYEEESPSRTIIQYIHDNYFLV
NLVDNDFPLDNCLWQVVEDTLELLNRPTQNARETEAP

MTHFD1 sequence (UniProt: P11586)

MAPAEILNGKEISAQIRARLKNQVTQLKEQVPGFTPRLAILQVGNRDDSNLYINVKLKAAEEIGIKATHIKLPRTTTESEVMKYITSLNED
STVHGFLVQLPLDSENSINTEEVINAIAPEKDVDGLTSINAGKLARGDLNDCFIPCTPKGCLELIKETGVPIAGRHAVVVGRSKIVGAPM
HDLLLWNNATVTTCHSKTAHLDEEVNKGDILVVATGQPEMVKGEWIKPGAIVIDCGINYVPDDKKPNGRKVVGDVAYDEAKERASF
ITPVPGGVGPMTVAMLMQSTVESAKRFLEKFKPGKWMIQYNNLNLKTPVPSDIDISRSCKPKPIGKLAREIGLLSEEVELYGETKAKVL
LSALERLKHRPDGKYVVVTGITPTPLGEGKSTTTIGLVQALGAHLYQNVFACVRQPSQGPTFGIKGGAAGGGYSQVIPMEEFNLHLTG
DIHAITAANNLVAAAIDARIFHELTQTDKALFNRLVPSVNGVRRFSDIQIRRLKRLGIEKTDPTTLTDEEINRFARLDIDPETITWQRVLD
TNDRFLRKITIGQAPTEKGHTRTAQFDISVASEIMAVLALTTSLEDMRERLGKMVVASSKKGEPVSAEDLGVSGALTVLMKDAIKPNL
MQTLEGTPVFVHAGPFANIAHGNSSIIADRIALKLVGPEGFVVTEAGFGADIGMEKFFNIKCRYSGLCPHVVVLVATVRALKMHGGG
PTVTAGLPLPKAYIQENLELVEKGFSNLKKQIENARMFGIPVVVAVNAFKTDTESELDLISRLSREHGAFDAVKCTHWAEGGKGALAL
AQAVQRAAQAPSSFQLLYDLKLPVEDKIRIIAQKIYGADDIELLPEAQHKAEVYTKQGFGNLPICMAKTHLSLSHNPEQKGVPTGFILPI
RDIRASVGAGFLYPLVGTMSTMPGLPTRPCFYDIDLDPETEQVNGLF

c

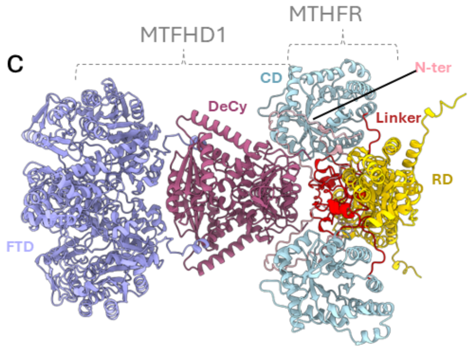


**Supplementary Fig. 6 | Follow-up analyses support interactions between MTHFR, MTHFD1, and GCN1.**

**a,** Immunoprecipitation of $MTHFR_{WT}$ and empty vector (EV) over-expressed in HEK293T MTHFR knock-out cells. Western blot using cell lysate prior to affinity pull down and after immunoprecipitation (IP) visualizing MTHFR with a primary antibody targeting the Flag-tag sequence. Corresponds to Figure 3a. **b,** Amino acid sequences of MTHFR (above) and MTHFD1 (below) used for Alphafold3 prediction in Figure 3c. **c,** Highest score AlphaFold 3 prediction of MTHFR and MTHFD1 without ligands. Colors and ligands as described in Figure 3.

**Supplementary Table 3 | Primers used in site directed mutagenesis, cloning and gene editing**

| | | |
|---|---|---|
| **Site directed mutagenesis of MTHFR_T34A** | MTHFR-T34A-F: | GATAGTTCGAGATGTTCCGCCCCGGGCCTGGAC |
| | MTHFR-T34A-R: | CTCGGGGTCCAGGCCCGGGGCGGAACATCTCGA |
| **LIC cloning of MTHFR_38-656** | LIC forward | TACTTCCAATCCATGGACCCTGAGCGGCATGAG |
| | LIC reverse | TATCCACCTTTACTGGATGGAGCCTCCGTTTCTCTC |
| **MTHFR-KO** | guide seq | GGGAGGCTTCAACTACGCAGTGG |
